# TFAP2 transcription factors are regulators of lipid droplet biogenesis

**DOI:** 10.1101/282376

**Authors:** Cameron C Scott, Stefania Vossio, Jacques Rougemont, Jean Gruenberg

## Abstract

How trafficking pathways and organelle abundance adapt in response to metabolic and physiological changes is still mysterious, although a few transcriptional regulators of organellar biogenesis have been identified in recent years. We previously found that the Wnt signaling directly controls lipid droplet formation, linking the cell storage capacity to the established functions of Wnt in development and differentiation. In the present paper, we report that Wnt-induced lipid droplet biogenesis does not depend on the canonical TCF/LEF transcription factors. Instead, we find that TFAP2 family members mediate the pro-lipid droplet signal induced by Wnt3a, leading to the notion that the TFAP2 transcription factor may function as a “master” regulator of lipid droplet biogenesis.

## Main Text

Cellular adaptation to a changing local environment is imperative for survival and proliferation. This is affected through a collection of sensing and signalling pathways that integrate information about the local environment and induce the requisite changes in various cellular programs that control organelle abundance and function, through multiple routes, including the modulation of transcription. In recent years, several transcriptional ‘master regulators’ of organellar biogenesis have been reported for mitochondria (Jornayvaz and Shulman, 2010), autophagosomes (Kang et al., 2012, Chauhan et al., 2013) and lysosomes (Sardiello et al., 2009). While the functional details of this control are still under investigation, coordinated transcriptional control of specific organelles is an emerging theme in cell biology.

Lipid droplets are primary storage organelle for neutral lipids in the cell (Meyers et al., 2017). Intriguingly, the number and nature of these organelles vary greatly, both over time within a cell, and between cell types (Thiam and Beller, 2017). While a major function of lipid droplets is clearly as the storehouse of triglycerides and sterol esters, the diversity and variation of this organelle likely reflect the number of reported alternate functions of lipid droplets such as regulation of inflammation, general metabolism, and host-pathogen interplay (Barisch and Soldati, 2017, Melo and Weller, 2016, Konige et al., 2014). Despite the recognized importance of this organelle in health and disease, little is known of the signalling systems or proximal transcriptional regulators that control lipid droplet biogenesis, function and turnover in cells.

Recently, we used genome-wide, high-content siRNA screens to identify genes that affect cellular lipids. This analysis revealed that the Wnt ligand can potently stimulate lipid droplet accumulation in multiple cell types through upstream elements of the Wnt signalling pathway, including the canonical surface receptors and adenomatous polyposis coli (APC), a component of the destruction complex (Scott et al., 2015). Therefore, to further characterize the signalling cascade leading to the accumulation of lipid droplets after Wnt addition, we tested the key components from the canonical Wnt signalling pathway for a role in lipid droplet regulation, starting with the core enzyme of the destruction complex, GSK3B (Fig.1A). We found that overexpression of both the wild-type, and the constitutively-active S9A (Stambolic and Woodgett, 1994) mutant of GSK3B were capable of attenuating lipid droplet accumulation in response to Wnt3a-treatment (Fig. 1B), consistent with the function of GSK3B activity as a negative regulator of Wnt signalling. Further, siRNAs to GSK3B were sufficient to induce lipid droplet accumulation (Fig. 1C) — much like we had shown after gene silencing of APC, another member of the destruction complex (Scott et al., 2015).

**Figure 1.**
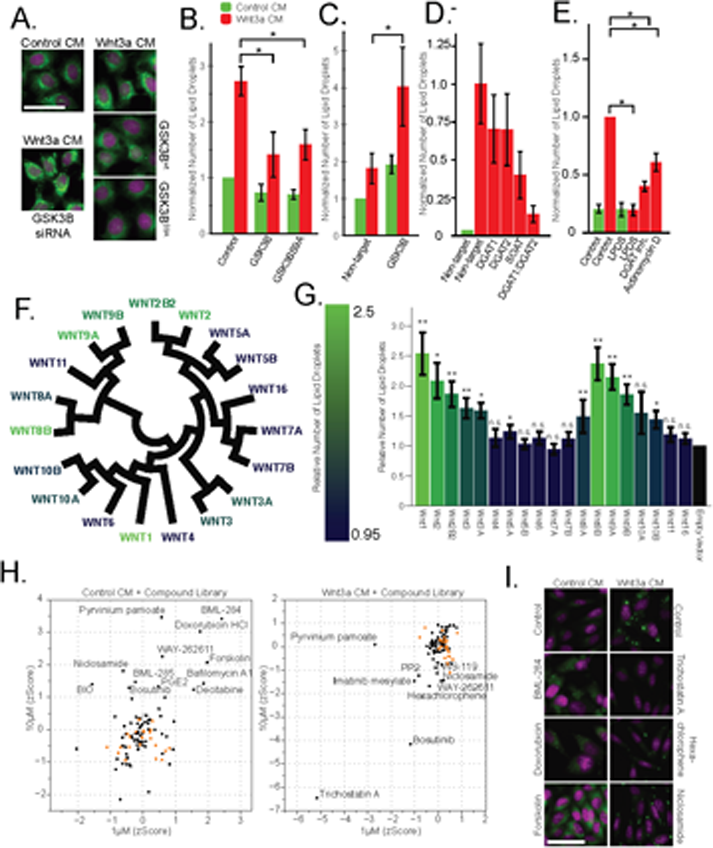
The Wnt pathway and the regulation of lipid droplets. **A.** HeLa-MZ cells were transfected with plasmids encoding wt or S9A mutant of GSK3B 24h before the addition of Wnt-3a- or control-conditioned media for a further 24h. Cells were fixed, labeled with BODIPY (lipid droplets, green) and Hoechst 33342 (nuclei, magenta), and imaged by light microscopy. **B.** The number of lipid droplets was quantified by automated microscopy (bar graph), and the data are presented as the mean number of lipid droplets per cell of 5 independent experiments ± SEM, normalized to the control condition. **C-D.** As in (A), except that HeLa-MZ cells (**C**) or L Cells (**D**) were transfected with siRNAs against the indicated targets for 48h, before the addition of Wnt3a. Efficient silencing was confirmed by qPCR (Fig. 1 - Fig.Sup. 1B) and the data are presented as the mean number of lipid droplets per cell of 3 independent experiments ± SEM, normalized to the control condition. **E.** L cells were incubated with the indicated compounds together with Wnt3a for 24h, processed and analyzed as in (A) and the data are presented as the mean number of lipid droplets per cell of 5 independent experiments ± SEM, normalized to the control condition. **F.** Evolutionary relationship of the 19 Wnt ligands. Colour indicates ability to induce lipid droplets as detailed in (G). **G.** L Cells were transfected with plasmids containing each of Wnt ligand for 48h, imaged and analyzed as in (A). Data are normalized to the empty vector control and were tested for significance (*, <0.05; **, <0.005, n.s., not significant) and are presented as the mean number of lipid droplets per cell of 2 independent replicates of the screen ± SEM, normalized to the control condition. The data are color-coded from a high (light) to a low (dark) number of lipid droplets induced by each Wnt ligand. **H-I.** High content image-based screen of a library of compounds that affect the Wnt pathway in HeLa-MZ cells. Cells were incubated for 24h with Wnt-3a- or control-conditioned media for 24h in the presence of the compounds at 1µM and 10µM, fixed, labeled with BODIPY (lipid droplets) and Hoechst 33342 (nuclei) and imaged by automated microscopy. The number of droplets per cell was counted and the zscores established, in order to quantify the ability of each compounds to induce lipid droplets in untreated cells (G, left panel), or to inhibit lipid droplet formation in Wnt3a-treated cells (G, right panel). Panel H illustrates the effects of compounds that induce droplet formation (left column) or that do (Trichostatin A) or do not (niclosamide, hexachlorophene) inhibit droplet formation (right column) in Wnt3a-treated cells. Nuclei are in magenta, and lipid droplets in green. Green bars, control-conditioned media; Red bars, Wnt3a-conditioned media. (*) indicates a p-value <0.05. See also Figure 1 – Figure Supplement 1, and Figure 1 – Source Data 1.

We next investigated the possible role of the downstream targets of the Wnt pathway at the transcription level. Surprisingly, siRNAs to TCF/LEF transcription factors relevant in canonical Wnt signalling, failed to affect lipid droplet accumulation - and yet there is no doubt that lipid droplet accumulation in response to Wnt3a is transcriptionally mediated. Indeed, the expression of SOAT1, DGAT1 and DGAT2, which encode key-enzymes of lipid droplet formation, increased in response to Wnt3a-treatment (Scott et al., 2015). In addition, silencing these genes inhibited lipid droplet accumulation in response to Wnt3a, most potently in combination with each other (Fig. 1D), or with specific inhibitors to DGAT1 or SOAT (Fig. 1E, Fig. 1 - Fig.Sup. 1A). Finally, Wnt3a-induced lipid droplets were decreased after treatment with the general inhibitor of transcription Actinomycin D (Fig. 1E). While these data altogether confirmed that the Wnt pathway was mediating the pro-lipid droplet signal, our inability to link lipid droplet induction to TCF/LEF led us to consider the possibility that a branching signalling path, under the control of GSK3B and/or ß-catenin but not the canonical Wnt transcription factors TCF/LEF, was inducing the accumulation of lipid droplets in cells. To explore this possibility, we initiated several parallel and complementary systems-level analyses with the aim to identify the transcriptional regulators proximal to lipid droplet biogenesis.

To better understand the nature of the pro-lipid droplet signal induced by Wnt, we revaluated the involvement of individual components of the Wnt signalling pathway in the induction of lipid droplet accumulation. First, we tested the ability of each of the 19 human Wnt ligands to induce lipid droplet accumulation in L Cells by transfection and autocrine or paracrine induction of the Wnt pathway (Fig. 1F-G). Wnt ligands displayed a broad, but not universal capacity to induce lipid droplet accumulation that paralleled both their evolutionary pedigree (Fig. 1F), and previously reported abilities to activate canonical Wnt signalling as measured by a TCF/LEF reporter system (Najdi et al., 2012). This confirmed that the pro-lipid droplet signal was indeed transiting, at least initially, through canonical Wnt signalling components.

To systematically assess the involvement of the remaining components of the Wnt pathway for involvement in the lipid droplet response, we performed a targeted screen for factors influencing lipid droplet accumulation using a library of 73 compounds selected for known interactions with elements of the Wnt pathway (see Methods). We tested the library in both, Wnt3a-stimulated conditions to assess any inhibitory activity of lipid droplet accumulation, and unstimulated conditions to identify compounds with the ability to induce the phenomenon (Fig. 1 – Source Data 1). Indeed, treatment with several compounds reported to activate the Wnt pathway induced lipid droplet accumulation, such as BML-284 (activator of ß-catenin (Liu et al., 2005)), doxorubicin (activator of Wnt signalling (Dai et al., 2009)), and forskolin (activation via PKA-Wnt crosstalk (Zhang et al., 2014)) (Fig. 1H-I). Conversely, known inhibitors of the pathway such as Tricostatin A (epigenetic regulator of DKK1 (Vibhakar et al., 2007)) hexachlorophene (ß-catenin inhibitor (Park et al., 2006)), and niclosamide (inducer of LRP6 degradation (Lu et al., 2011)) significantly decreased the appearance of lipid droplets in response to Wnt3a (Fig. 1H-I).

While these data certainly confirm the role of Wnts in regulating lipid droplets, they did not reveal the pathway linking the Wnt destruction complex to the transcriptional changes we observed (Fig. 1A-E, (Scott et al., 2015)). Given these results, and the large number of β-catenin-independent targets of the destruction complex (Kim et al., 2009), we initiated several strategies to identify candidate regulators, in particular the proximal transcription factors directly upstream from lipid droplet biogenesis. Our aim was to identify factors linked to Wnt signalling and to characterize the signalling pathway from ligand-stimulation to lipid droplet accumulation.

We first started by taking the subset of genes annotated as ‘transcription factor activity’ (GO:0000988) from our primary genome-wide siRNA screen data (Scott et al., 2015) to identify transcription factors that influenced cellular cholesterol levels in the cell (Fig. 2 - Fig.Sup. 1A; Fig. 2 – Source Data 1). While several of these candidates have well established roles in regulating general proliferation (i.e. MACC1, JDP2, SP3, TP53, ZNF217, TAF1), or links to the WNT pathway in keeping with our findings (i.e. GLI3, SIX2, SOX9, FOXK2, BARX1), we were particularly interested in identifying candidates with reported functions in lipid metabolism. The latter subset included ARNT2, STAT3, KDM3A, ATF5, KLF5, KLF6 and members of the TFAP2 (AP-2) transcription factor family (see below).

As a second approach to search for candidate transcriptional regulators of lipid droplets we compared existing transcriptome data of Wnt3a-treated cells with that of other conditions known to induce the accumulation of lipid droplets in tissue culture cells. It is well-established in the formation of lipid droplets is stimulated artificially by the addition of exogenous fatty acids, and the process has been studied at the transcriptional level in multiple studies. We therefore combined our Wnt3a gene array data (Scott et al., 2015) with three published datasets of mRNA levels after treatment of cells with fatty acids or knockout of lipid droplet regulatory factors (Li et al., 2010, Lockridge et al., 2008, Shaw et al., 2013). Our rationale was to identify the relevant transcription factors required for the induction of lipid droplet biogenesis by inferring from the expression data what is the common set of transcription factors active in response to Wnt3a and / or to the modulation of lipid droplets by fatty acid treatment or gene knockout. This analysis involved testing for over-representation of genes annotated to be regulated by a specific transcription factor in the set of the most perturbed genes after either treatment. Wnt3a-stimulation influenced a larger number of transcriptional regulators (172) as compared to lipid droplet modulation (91), but the vast majority of this subset (>75%) were also changed by Wnt3a (Fig. 2 - Fig.Sup. 1B).With this approach, we found that many candidate transcription factors known to function in both lipid metabolism and adipogenesis were influenced by Wnt3a-treament and fatty acid perturbation (Fig. 2 – Source Data 2), including TFAP2A (p-value Wnt3a treatment: 3.5 × 10^−9^; p-value fatty acid perturbation: 4.8 × 10^−4^).

As a third systems-level approach, we undertook a direct examination of the promotor regions of annotated lipid droplet proteins as was used by Sardiello and colleagues to identify the transcription factor TFEB and the CLEAR element as master regulators of lysosome biogenesis (Sardiello et al., 2009). We collected and examined the upstream promoter sequences for the 145 proteins annotated as ‘Lipid Droplet’ (GO:0005811; Fig. 2 – Source Data 3) and tested for over-represented sequence motifs. Among the most enriched motifs in the upstream promotor region of lipid droplet genes were motifs identified as TFAP2A (pValue: 9.0 × 10^−3^) and TFAP2C (pValue: 1.8 × 10^−5^) binding sites (Fig. 2 - Fig.Sup. 2A). Further analysis revealed that 74 of the ‘Lipid Droplet’ proteins contained at least one of the TFAP2A or TFAP2C annotated binding sites (Fig. 2 - Fig.Sup. 2B, Fig. 2 - Source Data 3) which were generally present within the first few hundred base-pairs from the start site (Fig. 2 - Fig.Sup. 2C).

The TFAP2 (AP-2) family of basic helix-span-helix transcription factors have been long recognized to play key roles during development. Yet, little else is known regarding their function in adult animals where these proteins are expressed, although various TFAP2 homologs have been linked to tumour progression in cancer models (Eckert et al., 2005). The family consists of five proteins in human and mouse, and are thought to form homo- and heterodimers that bind to similar promotor sequences albeit with different affinities. Given our observation that downstream genes known to be regulated by TFAP2 family members change in response to Wnt3a (Fig. 2 - Fig.Sup. 1B), that the silencing of TFAP2 genes can regulate cholesterol amounts in cells (Fig. 2 - Fig.Sup. 1A), and that TFAP2 consensus sites are over-enriched in the promoter sequences of genes encoding for lipid droplet proteins (Fig. 2 - Fig.Sup. 2A), we began to suspect that TFAP2 proteins mediate the pro-lipid droplet signal induced by Wnt3a. This notion was further buttressed by previous studies showing that TFAP2A directly interacts with both ß-catenin and APC (Li et al., 2009, Li et al., 2015) making this family of transcription factors our leading candidate for mediating the pro-lipid droplet signalling activity of Wnt3a.

To gain a detailed description of the transcriptional changes in the context of TFAP2, Wnt and lipid droplets, we performed an RNAseq determination of mRNA levels in cells treated with Wnt3a for short times (2h and 6h) with the aim to identify early factors of the transcriptional control relevant for lipid droplet biogenesis (Scott and Gruenberg, 2018). As expected, a pathway analysis of the most responsive genes at 2h found over-representation of terms related to mRNA processing, DNA binding and transcriptional regulation (Fig. 2A), consistent with the expected nature of the early Wnt3a-responsive genes. By 6h post-Wnt3a treatment, transcriptional regulators were still over-represented, but additional terms reflecting downstream effector pathways were present, including those related to glucose metabolism and endosomal trafficking, as well as fatty acid and cholesterol related genes (Fig. 2A) consistent with our previous findings (Scott et al., 2015). Indeed, the RNAseq analysis found that SREBF1 (Sterol Regulatory Element Binding Transcription Factor 1) mRNA levels were the most decreased of any transcription factor (0.60 of control) at the later time point (Fig. 2B).

**Figure 2.**
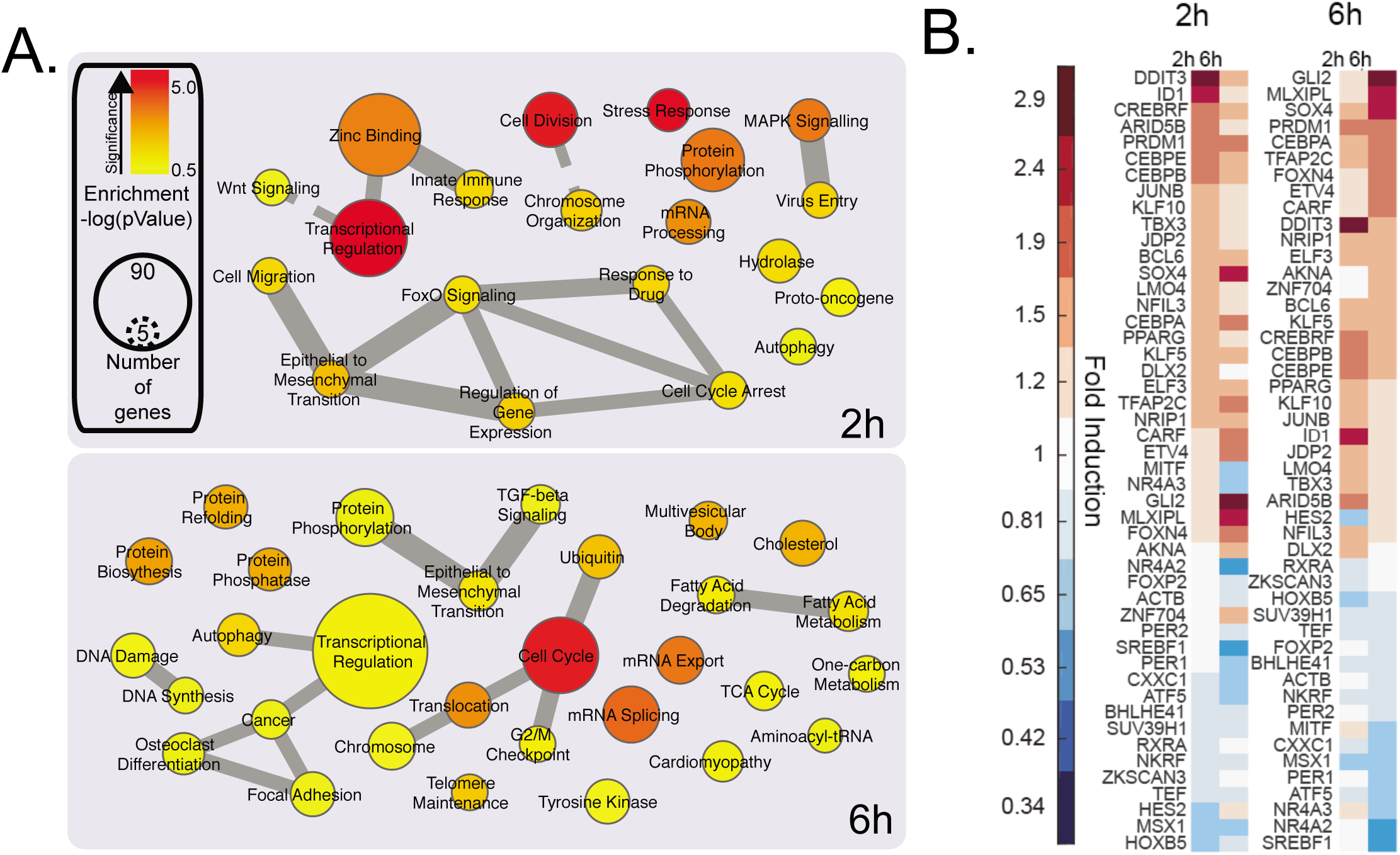
mRNA profiling and analysis of gene expression of cells treated with Wnt3a. A-B. HeLa-MZ cells were treated with control- or Wnt3a-conditioned media for 2h or 6h before RNA isolation and RNAseq analysis. Panel (A) shows the pathway enrichment of perturbed mRNAs. Node size indicates number of genes in each ontology and colour the statistical strength of the enrichment. Edge thickness indicates the strength of overlap of related ontologies. From (A), the perturbation of transcription factors amounts in response to Wnt3a is shown in panel B. See also Figure 2 – Figure Supplement 1 to 4, and Figure 2 – Source Data 1 to 3.

Intriguingly, the most upregulated transcription factor at early times was DDIT3 (DNA Damage Inducible Transcript 3, also known as CHOP) — and increased DDIT3 could readily be detected at the mRNA and protein level (Fig. 2 - Fig.Sup. 3A-B). This member of the CCAAT/enhancer-binding (CEBP) protein family is a potent and direct inhibitor of SREBF1 transcription (Chikka et al., 2013) and a known regulator of cellular lipid metabolism. While this interaction may contribute to the decrease in cellular free cholesterol and the downregulation of cholesterol metabolic enzymes after Wnt3a treatment (Fig. 2A, (Scott et al., 2015)), overexpression of constitutively-active SREBF1 truncations (Shimano et al., 1997) had essentially no effect on the formation of lipid droplets, whether Wnt3a was present or not (Fig. 2 - Fig.Sup. 4A). Neither did treatment with the S1P/SREBF1 inhibitor PF- 429242 (Hawkins et al., 2008) (Fig. 2 - Fig.Sup. 4B) in either Wnt3a-stimulated, or naïve conditions. These observations suggest that while relevant to the observed changes in cellular cholesterol homeostasis generated by Wnt3a, putative DDIT3-induced changes in SREBF1 expression have no significant role in lipid droplet accumulation in response to Wnt3a.

Along with DDIT3 and SREBF1, our list of early Wnt3a-responsive transcription factors includes several that have known functions in regulating cellular lipid homeostasis such as CEBPB and CEBPE, KLF5, KLF10, PPARG, MLXIPL, and PER2. Our list also included the TFAP2 family member TFAP2C, which our datamining strategies had already identified as involved in lipid homeostasis, and as a candidate transcription factor controlling lipid droplet biogenesis. In fact, 6h post-Wnt3a TFAP2C (1.72-fold) was among the most upregulated transcription factors (Fig. 2B).

Given that our datamining efforts identified TFAP2 family members as putative transcription factors regulating lipid droplet proteins and that TFAP2C was among the most upregulated transcription factors in response to Wnt3a (Fig. 2B), we next investigated whether TFAP2 family members played a direct role in regulating lipid droplets. To this end, we tested whether Wnt3a retained the ability to induce lipid droplets after TFAP2 depletion by RNAi. While knock-down of either TFAP2A or TFAP2C had no or only a modest effect, tandem silencing of both homologs produced a marked reduction in the number of lipid droplets present in cells in response to Wnt3a (Fig. 3A-B). In keeping with this finding, these siRNA treatments diminished the mRNA levels of SOAT1, a key enzyme proximal to the production of lipid droplets (Fig 3C) that mediate the production of cholesteryl esters (Chang et al., 2001).

**Figure 3.**
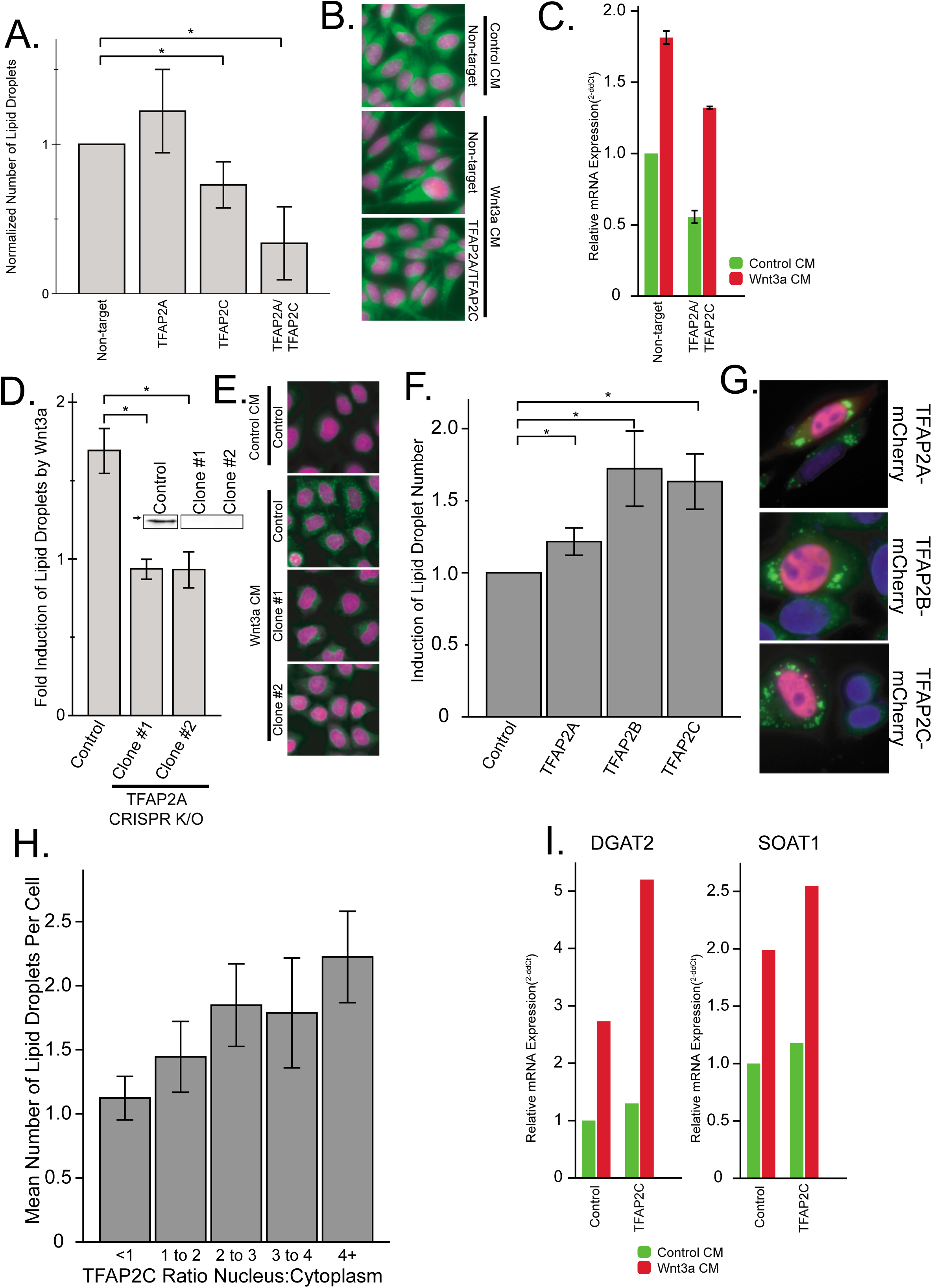
The TFAP2 family of transcription factors are both necessary and sufficient to mediated lipid droplet accumulation. **A-B.** L Cells were treated with siRNA against the indicated targets for 48h before the addition of Wnt3a-conditioned medium for an additional 24h. Cells were then fixed, labeled, imaged and analyzed by automated microscopy as in Fig 1A. In (A), data are presented as the normalized mean number of lipid droplets per cell of 5 independent experiments ± SEM. (*) indicates a p-value <0.05. Cells treated with non-target siRNAs or with siRNAs to both TFAP2A and TFAB2C are shown in panel B (nuclei in magenta; lipid droplets in green). **C.** L cells were treated with siRNAs against the indicated targets as in (A), before the addition of Wnt3a- or control-conditioned media for an additional 24h. RNA was isolated and analyzed by qPCR, and that data are expressed relative to the non-target control and are presented as the mean number of lipid droplets per cell of 2 independent experiments ± SEM. **D-E.** HeLa-MZ cells were transfected with targeted CRISPR/Cas9 plasmids against TFAP2A. The corresponding knock-out clones as well as control cells were treated with Wnt3a conditioned-media for 24h. In **D**, the number of lipid droplets was quantified as in Fig 1A and is expressed as fold induction relative to the control cells in 5 independent experiments ± SEM. Inset: TFAP2A protein levels of each clone determined by Western blot. Arrow indicate position of 50 kDa marker. Representative images are shown in E (<u>nuclei in magenta; lipid droplets in green</u>). **F-H.** L Cells were transfected or not with mCherry-tagged TFAP2 family members for 48h before fixation, labeling and imaging as in Fig 1A. The mean number of lipid droplets per cell expressing each mCherry-tagged TFAP2 protein was counted, and is expressed, as in panel (D), as fold induction relative to the control cells in 6 independent experiments ± SEM. (*) indicates a p-value <0.05. Panel G shows cells expressing each mCherry-tagged TFAP2 protein (Blue, nucleus; Green, lipid droplets; Red, TFAP2-mCherry fusion proteins), and panel H shows the number of lipid droplets per cell in cells overexpression TFAP2C-mCherry, binned by their nuclear:cytoplasmic distribution. Data are the mean lipid droplets per cell ± SEM for 450 cells. **I.** L Cells were treated as in **F** before extraction and determination of the indicated mRNAs by qPCR. Data are representative of two independent experiments.

As an alternative approach, we used CRISPR/Cas9 gene knockout to generate HeLa-MZ cells clones lacking TFAP2A (Fig. 3D-E). While tandem depletion by RNAi was necessary to reduce lipid droplet production after Wnt3a addition in L Cells, two knockout clones of TFAP2A demonstrated a complete lack of change in lipid droplet number after Wnt3a stimulation (Fig. 3D-E), indicating that TFAP2 family members exhibit complementary functions. Together, these results imply that TFAP2A/TFAP2C are necessary for mediating the pro-lipid droplet signal of the Wnt pathway.

We next sought to determine if the TFAP2 family is sufficient for the induction of lipid droplets. For this, we fused full-length TFAP2A, TFAP2B, and TFAP2C to mCherry and overexpressed these as exogenous chimeras. As transcription factors the TFAP2 family members function in the nucleus, the mCherry-tagged TFAP family members exhibited a somewhat heterogeneous distribution between cytoplasm and nuclei, presumably because of variations in expression levels. Impressively, the number of lipid droplets per cell increased with increased nuclear localization of each TFAP2 family member (Fig. 3H), and, in cells with TFAP2 nuclear localization, the expression of each family member was clearly sufficient to cause lipid droplet biogenesis (Fig. 3F-G). Moreover, the expression of TFAP2C was also able to trigger the expression of lipid droplet enzymes (Fig. 3I), further supporting the notion that TFAP2 family members function as transcriptional regulators of lipid droplet biogenesis.

In totality, these findings demonstrate TFAP2A, TFAP2B and TFAP2C are sufficient to induce biogenesis of lipid droplets when expressed in cells, and members of the TFAP2 family are required to mediate the accumulation of lipid droplets seen in response to Wnt-stimulation. This finding is in keeping with a previous report that targeted overexpression of TFAP2C in mouse liver induced steatosis, accumulation of fat, and eventual liver failure (Holl et al., 2011), and given the enrichment in TFAP2 binding sites in lipid droplets proteins (Fig. 2 - Fig.Sup. 2), implicate the TFAP2 family as central regulators of lipid droplet biogenesis.

Given our observation that TFAP2C expression correlates with DDIT3 expression in cells after Wnt3a treatment (Fig. 2B), we investigated the role of DDIT3 in control of lipid droplets. Indeed, while changes in SREBF1 expression induced by DDIT3 do not seem to be involved in lipid droplet accumulation (Fig. 2 - Fig.Sup. 4), this process may be regulated by DDIT3 itself. We started by examining the promotor region of the transcription factors whose mRNA levels were impacted by Wnt3a-treatment (Fig. 2B) for reported DDIT3::CEBPA consensus sites (Ubeda et al., 1996). Intriguingly, not only did we find such a site upstream of the TFAP2C gene, but there was a DDIT3::CEBPA consensus site in 7 out of the 10 lipid-related transcription factors identified in our RNAseq (Fig. 4 - Fig.Sup. 1A), a 1.8-fold enrichment over the frequency in the total Wnt3a-influenced transcription factor set. This lends further support for the view of DDIT3 as an important transcriptional regulator of lipid homeostasis in cells (Chikka et al., 2013).

To address this notion directly, we next tested whether DDIT3 itself was able to influence both TFAP2 family members, and the number of lipid droplets in cells. Indeed, overexpression of DDIT3-mCherry chimera resulted in an increase in the mRNA levels of lipid droplet enzymes (Fig. 4A). Further, silencing of DDIT3 both diminished SOAT1 mRNA levels (Fig. 4B) and lipid droplet accumulation in response to Wnt3a (Fig. 4C). These observations suggest that DDIT3 plays a regulatory role in lipid droplet biogenesis, presumably in concert with TFAP2.

**Figure 4.**
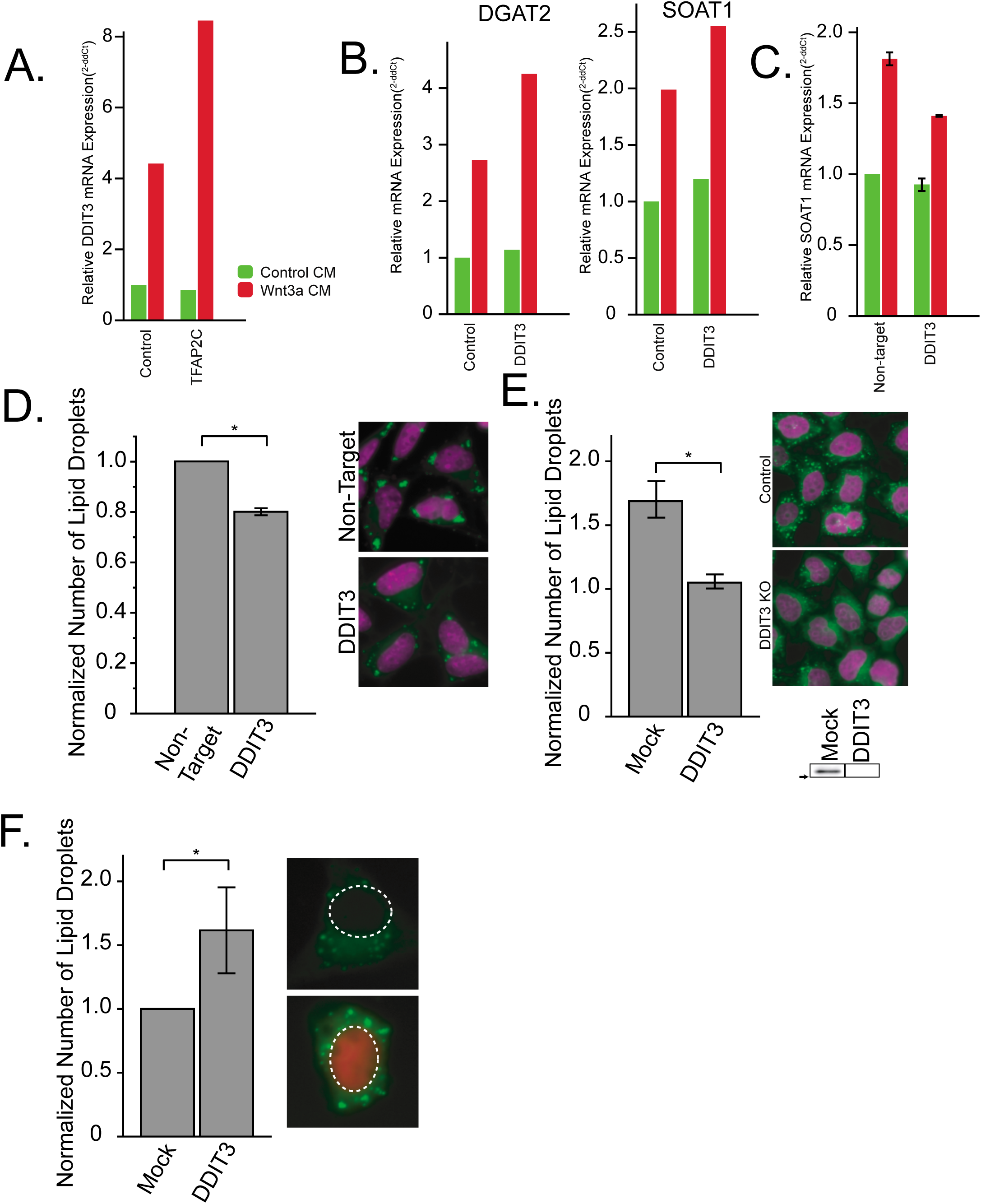
DDIT3 is both necessary and sufficient to mediated lipid droplet accumulation. **A-B.** qPCR of DDIT3, DGAT2 and SOAT1 after overexpression of TFAP2C-mCherry (A), DDIT3-mCherry (B), or siRNAs to DDIT3 (C). Data are representative of two independent experiments. **C.** L cells were treated, processed and analyzed like in Fig 3A, except that they were transfected with siRNAs to DDIT3 before stimulation with Wnt3a. Data are presented as the normalized mean number of lipid droplets per cell of 3 independent experiments ± SEM. (*) indicates a p-value <0.05. Panel E shows cells treated with non-target or anti-DDIT3 siRNAs (magenta, nucleus; green, lipid droplets). **D.** HeLa-MZ CRISPR/Cas9 clones were prepared and analysed as in Fig 3D. Data are expressed as fold induction relative to the control cells in 5 independent experiments ± SEM. Inset: DDIT3 protein levels of each clone determined by Western blot. Arrow indicate position of 25 kDa marker. Representative images are shown (nuclei in magenta; lipid droplets in green). **E.** L cells were treated, processed and analyzed like in Fig 3F, except that they were transfected with a plasmid encoding DDIT3-mCherry. Data are presented as the normalized mean number of lipid droplets per cell of 6 independent experiments ± SEM. (*) indicates a p-value <0.05. Panel E shows cells expressing or not DDIT3-mCherry (blue, nucleus; green, lipid droplets, red, DDIT3-mCherry fusion protein). See also Figure 4 – Figure Supplement 1.

We next tested the requirement of DDIT3 for the lipid droplet response directly by interfering with DDIT3 expression by both RNAi and CRISPR/Cas9. Silencing of DDIT3 expression diminished accumulation of lipid droplets both in response to Wnt3a stimulation (Fig. 4C; Fig. 4 - Fig.Sup. 1B), and after silencing of APC (Fig. 4 - Fig.Sup. 1B), an alternative strategy previously shown to be sufficient to induce lipid droplets (Scott et al., 2015). Further, knock-out clones lacking DDIT3 were completely non-responsive to Wnt3a with regards to lipid droplet number. In the context of the significant increase in DDIT3 message and protein (Fig. 2 - Fig.Sup. 3), these results suggest that induction of DDIT3 transcription in response to Wnt3a is necessary for lipid droplet biogenesis.

Given that the presence of DDIT3 was necessary to convey the pro-lipid droplet signal, we next tested whether over-expression of the transcription factor is sufficient to induce lipid droplet accumulation. As with TFAP2, overexpression of mCherry-tagged fusions of DDIT3 was sufficient to increase lipid droplet numbers (Fig. 4E; Fig. 4 - Fig.Sup. 1C) in transfected cells as compared to the control. In total, these data suggest that in addition to regulation of cholesterol metabolism (Chikka et al., 2013), DDIT3 may function as a more global regulator of cellular lipid homeostasis in part through regulation of the TFAP2 family of transcription factors.

## CONCLUSIONS

In conclusion, our data show TFAP2 family members function to modulate expression of lipid droplet proteins and induce the accumulation of lipid droplets in cells. We found TFAP2C is necessary to potentiate the pro-lipid droplet signal induced by Wnt3a, and expression of TFAP2 family members is sufficient to induce lipid droplet accumulation in cells (Fig. 3).

Not only do these data support the view that the TFAP2 family of transcription factors can function as regulators of lipid droplet biogenesis, they provide insight into the transcriptional network directing changes in lipid homeostasis and the accumulation of lipid droplets in response to Wnt stimulation. Our observations indicate that, in addition to the canonical TCF/LEF transcriptional response, Wnt signalling via APC, GSK3, and ß-Catenin induces transcription-mediated lipid changes. These include decreased levels of SREBF1, which is likely to contribute to the observed reduction in total membrane cholesterol levels (Scott et al., 2015), and increased expression of DDIT3 and TFAP2, which triggers lipid droplet biogenesis with storage of cholesterol esters and triglycerides. These observations also lead to the notion that via TFAP2 transcription factors, Wnt exerts pro-proliferative effects in developmental and in pathological contexts by leading to the accumulation of lipid droplets.

In addition, Wnt3a has been recently identified as an intra-cell synchronizer of circadian rhythms in the gut, controlling cell-cycle progression, and the mRNA of Wnt3a itself exhibits circadian oscillations under the control of the master clock regulators Bmal1 and Per (Matsu-Ura et al., 2016). Given this function, it is tempting to speculate that Wnts serve a more general role as a mid- or long-range circadian signalling intermediates, linking lipid metabolism to the master transcriptional clocks that coordinate metabolic functions with the day-night cycle. Because of its circadian nature, and capacity to induce lipid droplet accumulation in a broad range of cells types, Wnts could serve as a fundamental signal for cells to store lipids during times when nutrients are expected to be in excess for later use during periods of rest or fasting.

Both lipid droplets enzymes (Solt et al., 2012), and the volume of lipid droplets themselves (Uchiyama and Asari, 1984), vary in a circadian fashion, which is completely consistent with a role as neutral lipid storage sites. TFAP2 binding sites are over-represented in the promotors of circadian-controlled genes (Bozek et al., 2009), suggesting that the Wnt/TFAP2 control of lipid droplets is one mechanism by which the daily storage of lipids in lipid droplets, and cellular lipid homeostasis itself, is coordinated. Further, targeted overexpression of TFAP2C induced the equivalent phenotype of steatosis in mouse liver (Holl et al., 2011), underscoring the role of TFAP2 proteins in the regulation of neutral lipid metabolism.

The Wnt pathway is also not the first developmental signalling pathway found to exert key regulatory functions in directing energy metabolism. The FOXA family of transcription factors, fundamental for early embryogenesis, play core roles in directing glucose metabolism (Friedman and Kaestner, 2006), while SOX17 has been shown to have additional, non-developmental functions regulating lipid metabolism in the liver of adult animals (Rommelaere et al., 2014). Given the overlapping requirement for a molecular queue to synchronize energy storage during both embryo development and daily metabolic activity, it is not surprising that some of these systems have evolved to serve dual roles in both biological contexts. These findings support the view that both Wnt signalling, and the TFAP2 family of transcription factors have important (and possibly linked) roles in development and circadian lipid metabolism of the cell.

## Materials and Methods

### Cells, media, reagents and antibodies

HeLa-MZ cells, a line of HeLa cells selected to be amiable to imaging, were provided by Prof. Lucas Pelkmans (University of Zurich). HeLa cells are not on the list of commonly misidentified cell lines maintained by the International Cell Line Authentication Committee. Our HeLa-MZ cells were authenticated by Microsynth (Balgach, Switzerland), which revealed 100% identity to the DNA profile of the cell line HeLa (ATCC: CCL-2) and 100% identity over all 15 autosomal STRs to the Microsynth’s reference DNA profile of HeLa. L cells (ATCC: CRL-2648) and L Wnt3A cells (ATCC: CRL-2647) were generously provided by Prof. Gisou van der Goot (École Polytechnique Fédérale de Lausanne; EPFL) and cultured as per ATCC recommendations. Cells are mycoplasma negative as tested by GATC Biotech (Konstanz, Germany). Wnt3A-conditioned, and control-conditioned media was prepared from these cells by pooling two subsequent collections of 24h each from confluent cells.

Reagents were sourced as follows: Hoechst 33342 and BODIPY 493/503 from Molecular Probes (Eugene, OR); The DGAT1 inhibitor A-922500 and the Membrane Bound Transcription Factor Peptidase, Site 1 (S1P/SREBF) inhibitor PF-429242, and the mTOR inhibitor Torin-2 were from Tocris Bioscience (Zug, Switzerland); lipoprotein-depleted serum (LPDS) was prepared as previously described (Havel et al., 1955); anti-DDIT3 antibodies were from Cell Signaling (L63F7; Leiden, The Netherlands); anti-TFAP2A antibodies were from Abcam (ab52222; Cambridge, UK); fluorescently labeled secondary antibodies from Jackson ImmunoResearch Laboratories (West Grove, PA); 96-well Falcon imaging plates (#353219) were from Corning (Corning, NY); oligonucleotides and small interfering RNA (siRNA) from Dharmacon (SmartPool; Lafayette, CO) or QIAGEN (Venlo, The Netherlands) and the siRNAs used in this work were: GSK3B (S100300335); DGAT1 (S100978278); DGAT2 (S100978278); SOAT1 (S101428924); TFAP2A (J-062799); TFAP2C (J-048594); DDIT3 (J-062068); APC (S102757251).

Other chemicals and reagents were obtained from Sigma-Aldrich (St. Louis, MO). Transfections of cDNA and siRNAs were performed using Lipofectamine LTX and Lipofectamine RNAiMax (Invitrogen; Basel, Switzerland) respectively using the supplier’s instructions.

Plasmids encoding GSK3B and GSK3B^S9A^, and the SREBF truncations were from Addgene (Cambridge, MA); TFAP2A, TFAP2B, and DDIT3 coding sequences were obtained from the Gene Expression Core Facility at the EPFL, and TFAP2C from DNASU (Arizona State University) and cloned into appropriate mammalian expression vectors using Gateway Cloning (Thermo Fisher Scientific). Knock-out cell lines of TFAP2A and DDIT3 were obtained by clonal isolation after transfection of a pX330 Cas9 plasmid (Cong et al., 2013) with appropriate guide sequences (see Supplementary Methods).

### Lipid Droplet Quantitation

L Cells were seeded (6 000 cells/well) in imaging plates the day before addition of control-conditioned, or Wnt3a-conditioned media for 24h, before fixation with 3% paraformaldehyde for 20 min. Where necessary, cells were treated with siRNAs and Lipofectamine RNAiMax Reagent (Thermo Fisher) per the manufacturer’s instructions for 48 hours before seeding into imaging plates. Lipid droplets and nuclei were labelled with 1µg/mL BODIPY 493/503 (Invitrogen; D3922) and 2µg/mL Hoechst 33342 (Invitrogen; H3570) for 30 min, wash with PBS, and the plate sealed and imaged with ImageXpress Micro XLS (Molecular Devices, Sunnyvale, CA) automated microscope using a 60X air objective. Images were segmented using CellProfiler (Carpenter et al., 2006) to identify and quantify nuclei, cells, and lipid droplets.

### Wnt Ligand Screen

L Cells cells were seeded into image plates (6 000 cells/well) the morning before transfection with Lipofectamine 3000 (Invitrogen) as per the manufacturer’s instructions with a subset of untagged Wnts from the open source Wnt Project plasmid library (Najdi et al., 2012). Cells were fixed with 3% PFA after 48h, and lipid droplets quantified as above. Data were normalized to the number of lipid droplets in the empty vector condition. The circular phylogenetic tree was constructed using human Wnt sequences and the Lasergene (v12.1; DNAStar, Madison WI) bioinformatics software.

### Wnt Compound Screen

L Cells were seeded into imaging plates (6 000 cells/well) the day before addition of the Wnt Pathway Library (BML-2838; Enzo Life Sciences, Farmingdale, NY) at either 1µM or 10µM, with either control-conditioned or Wnt3a-conditioned media for 24h, before fixation and lipid droplet quantitation as above. Data were normalized by z-score and the average of quadruplicates.

### mRNA Determination

RT-PCR was carried out essentially as described (Brankatschk et al., 2012, Scott et al., 2015), after total RNA extraction using TRIzol Reagent (Life Technologies AG; Basel, Switzerland) according to manufacturer’s recommendation from monolayers of HeLa-MZ or L cells.

### Western Blotting

For Western blotting, cells were lysed with RIPA buffer plus protease inhibitors (1mM phenylmethylsulfonyl fluoride, 10µg/ml aprotinin, 1µM pepstatin, 10µM leupeptin) for 20 min. The lysates were sonicated, clarified by centrifugation at (14 000 RCF for 10 min at 4°C) and were separated by SDS-PAGE and blotted on nitrocellulose or polyvinylidene fluoride membranes.

### RNAseq

HeLa-MZ cells were treated in triplicate with control-conditioned, or Wnt3a-conditioned media for 2h or 6h before cells were collected and RNA isolated. Purification of total RNA was done with the RNeasy Mini Kit from Qiagen according to the manufacturer’s instructions. The concentration, purity, and integrity of the RNA were measured with the Picodrop Microliter UV-Vis Spectrophotometer (Picodrop), and the Agilent 2100 Bioanalyzer (Agilent Technologies), together with the RNA 6000 Series II Nano Kit (Agilent) according to the manufacturer’s instructions. Purified RNAs were sequenced with a HiSeq 4000 (Illumina). The datasets are available at EMBL-EBI ArrayExpress (https://www.ebi.ac.uk/arrayexpress/)under accession number E-MTAB-6623.

Sequencing reads were mapped to the hg38 genome using bowtie2 in local alignment mode. Then reads were attributed to known exons as defined by the ensembl annotation, and transcript-level read counts were inferred as described in (David et al., 2014). Differential expression was then evaluated by LIMMA (Law et al., 2014) using the log of rpkm values. The fold induction was determined as a ratio of mRNA amounts of Wnt3a to control. Genes with message levels increase more than 1.5-fold, or decreased less than 0.8-fold were collected and tested for pathway enrichment using DAVID bioinformatics resources (v6.8) (Huang da et al., 2009) and the resulting data compiled and plotted using Cytoscape (Shannon et al., 2003) and previously described Matlab scripts (Mercer et al., 2012).

### Wnt siRNA Screen Analysis

Data from the genome-wide siRNA screen (Scott et al., 2015) was used to produce the subset of annotated transcription factors (GO:0003700) cellular cholesterol levels after gene silencing provided by AmiGO 2 (version 2.4.26) (Carbon et al., 2009). Data were filtered for genes that increased or decreased total cellular cholesterol a z-score of 1.5 or greater.

### Transcription Factor Enrichment

Transcript data from cells treated with Wnt3a (E-MTAB-2872 (Scott et al., 2015)), or fatty acids (GSE21023 (Lockridge et al., 2008), GSE22693 (Xu et al., 2015), GSE42220 (Shaw et al., 2013)) were collected and mRNAs that significantly changed in response to treatment were compiled and analyzed for transcription factor enrichment using GeneGo (MetaCore) bioinformatics software (Thomson Reuters). Data from both conditions was compared and ordered by a sum of ranking.

### Promotor Analysis

Promotor sequences from the current (GRCh38) human genome were collected using Regulatory Sequence Analysis Tools (RSAT) software (Medina-Rivera et al., 2015) to collect the non-overlapping upstream promotor sequences (<3 000 bp) of genes of interest which were tested for enriched sequences using either the RSAT’s oligo-analysis or Analysis of Motif Enrichment (AME) module of MEME Suite (v.4.12.0) (McLeay and Bailey, 2010).

Genes with no non-overlapping upstream region were discarded from the analysis. Identification and quantitation of consensus binding sites was done with either RSAT’s matrix-scan or MEME Suite’s Motif Alignment and Search Tool (MAST) module using a cut-off of <0.0001 to define a consensus site. Consensus sites from JASPAR (Mathelier et al., 2016) were: DDIT3:CEBPA (MA0019.1), TFAP2A (MA0003.2, MA0810.1), TFAP2C (MA0524.1, MA0524.2).

### Construction of Gene Edited Knock Out Cell Lines

The guide sequences used to construct specific CRISPR/Cas9 vectors were determined using the CRISPR Design Tool(Ran et al., 2013) and were:

TFAP2A

Fwd: CACCGGAGTAAGGATCTTGCGACT

Rev: AAACAGTCGCAAGATCCTTACTCC

DDIT3

Fwd: CACCGGCACCTATATCTCATCCCC

Rev: AAACGACTGATCCAACTGCAGAGAC

These sequences were used to create insert the target sequence into the pX330 vector using Golden Gate Assembly (New England Biolabs) and transfected into cells as described in the Methods. Knock-out clones were isolated by serial dilution and confirmed by Western blotting and activity assays.

## Acknowledgements

We are grateful to Christian Iseli, Nicolas Guex, and Ioannis Xenarios from Vital-IT and the Swiss Institute of Bioinformatics for indispensable guidance on the interpretation of the transcriptomics data and critical reading of the manuscript. RNA was sequenced at the iGE3 Genomics Platform. of the University of Geneva (https://ige3.genomics.unige.ch). Support was from the Swiss National Science Foundation, the NCCR in Chemical Biology and LipidX from the Swiss SystemsX.ch Initiative, evaluated by the Swiss National Science Foundation (to J. G.). C. C. S. has been supported by fellowships from the Human Frontier Science Program and the Canadian Institutes of Health Research.

## Author Contributions

CCS carried out most of the experiments and analyses and wrote the manuscript; SV contributed the mRNA determinations and assisted with lipid droplet quantitation; JR contributed to the analysis of the RNAseq data; and JG supervised the entire project and wrote the manuscript.

## Competing Interests

The authors declare no competing interests.

## Materials & Correspondence

Correspondence and requests to Jean Gruenberg.

## Figure Source Data

Figure 1 – Source Data 1: Effect of Wnt pathway related compounds on lipid droplet induction.

Figure 2 – Source Data 1: Effect of silencing transcription factors on cellular cholesterol amounts.

Figure 2 – Source Data 2: Comparative enrichment of transcriptional targets in cells treated with Wnt3a or fatty acid perturbation.

Figure 2 – Source Data 3: TFAP2 family member consensus binding sites in lipid droplet genes.

---

**Figure 1 – Figure Supplement 1.**
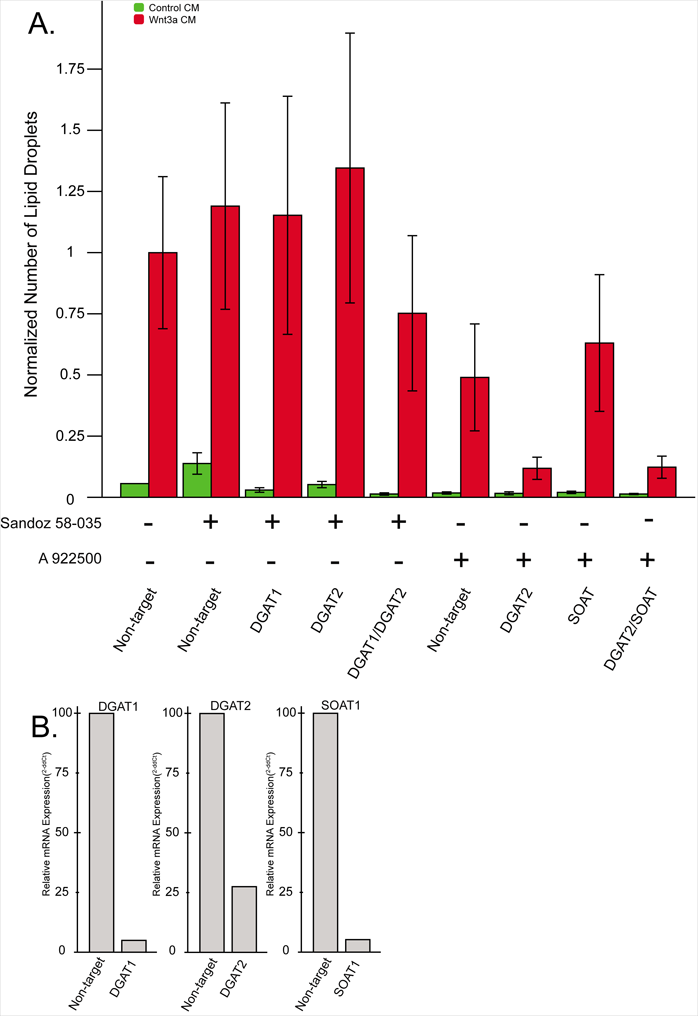
Lipid droplet accumulation in response to Wnt3a: combinatorial treatments against lipid droplet enzymes by RNAi and chemical inhibitors. **A.** L cells were treated with the indicated siRNAs for 48h before addition of the indicated compounds together with control, or Wnt3a, conditioned media and incubation for an additional 24h. Cells were then fixed, stained and lipid droplet number quantified as in Fig 1A. Data are presented as the normalized mean number of lipid droplets per cell of 3 independent experiments ± SEM. **B.** L Cells were treated as in Fig. 1A with the indicated siRNAs and the amounts of the corresponding mRNA were determined by qPCR.

**Figure 2 – Figure Supplement 1.**
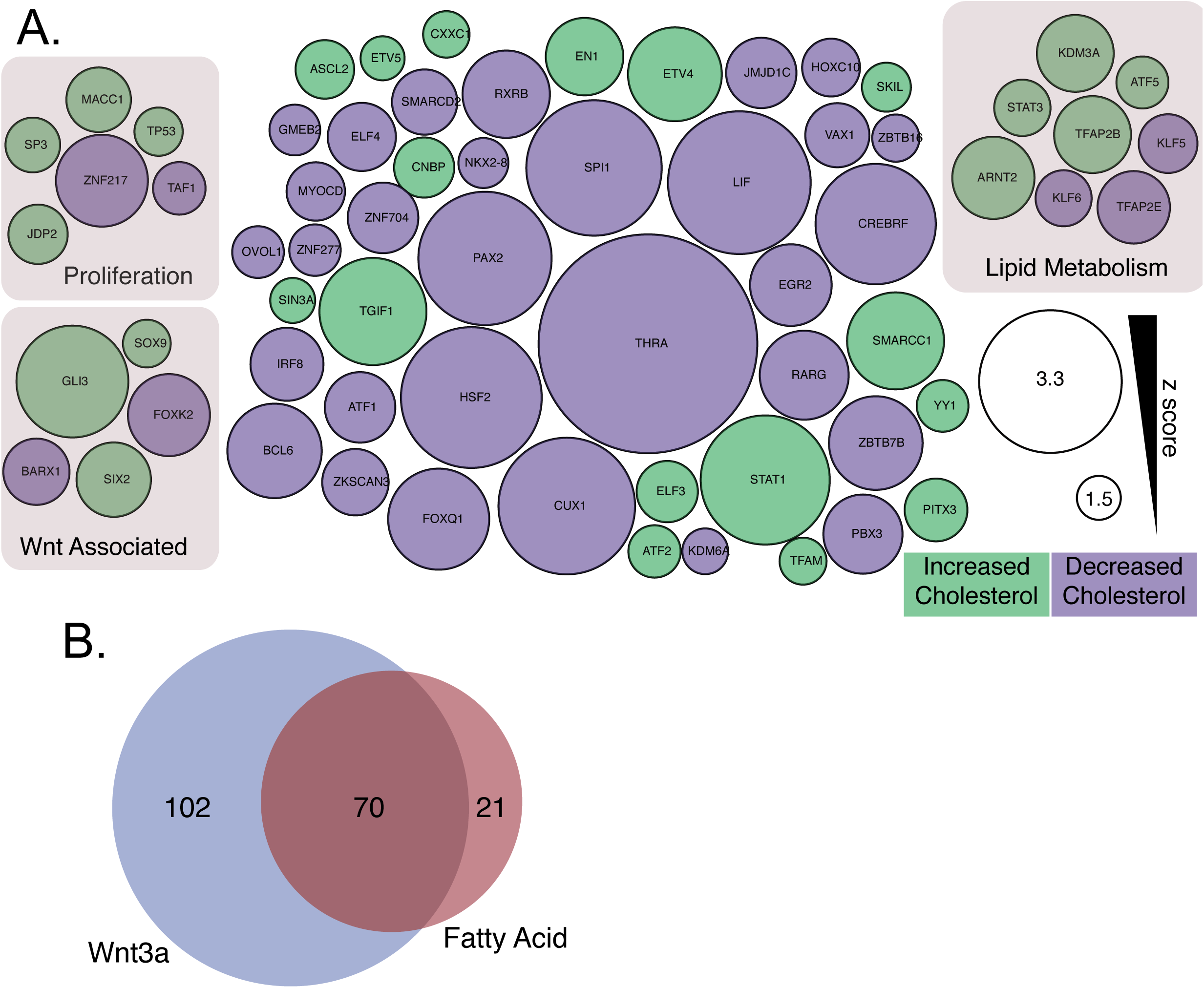
Datamining for putative lipid droplet transcriptional regulators. **A.** Transcription factors that influence cellular cholesterol amounts from a genome-wide screen of cholesterol regulatory genes (Scott et al., 2015). Node size is proportional to the absolute z-score difference from the control, and the colour indicates increased (green) or decreased (blue) cellular cholesterol levels. **B.** Existing mRNA profiling experiments of cells treated with Wnt3a (blue), or perturbations likely to induce lipid droplet accumulation (see methods) (pink), were analyzed for enrichment of genes linked to known transcriptions factors to identify candidate transcription factors linking Wnt3a to lipid droplet biogenesis.

**Figure 2 – Figure Supplement 2.**
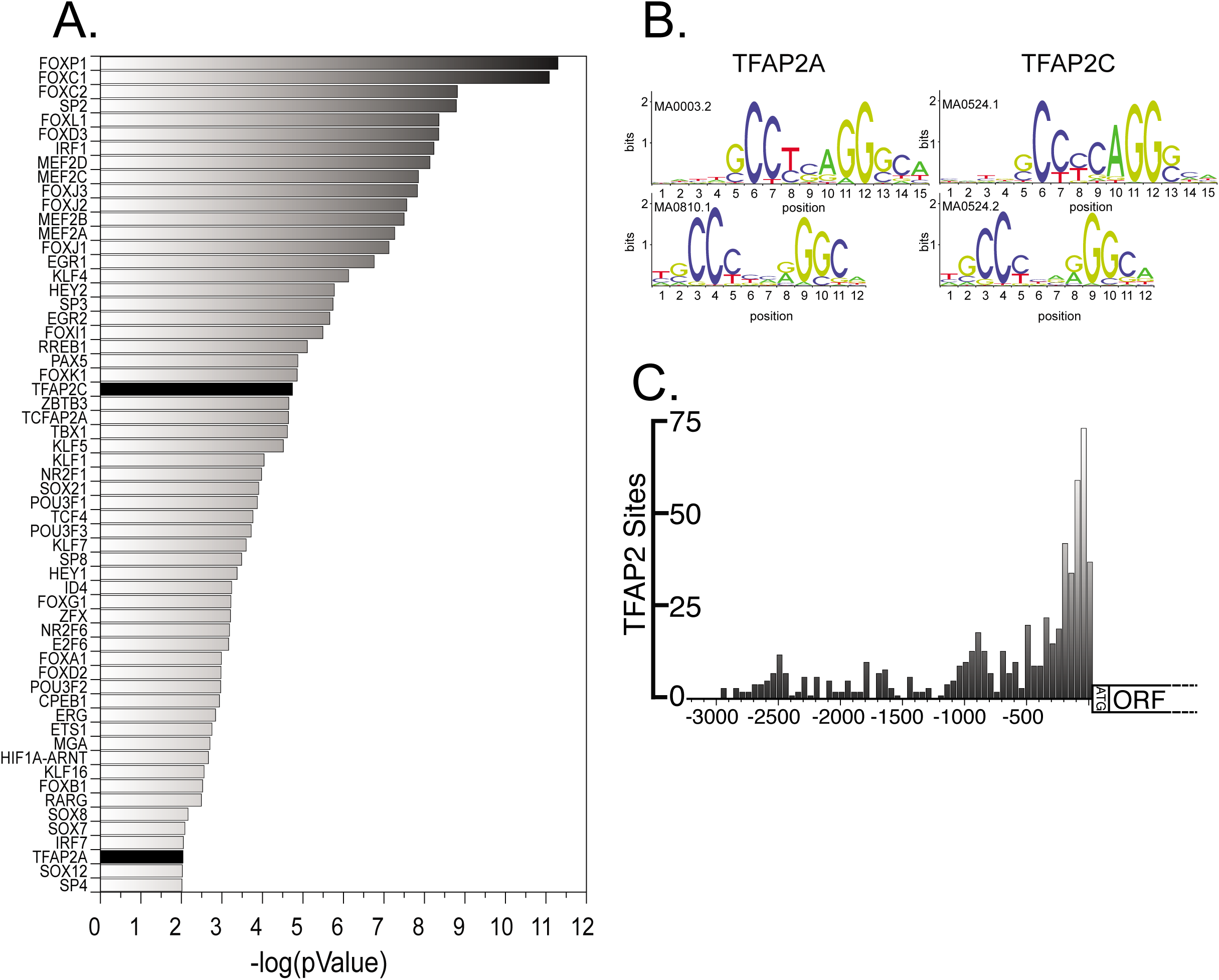
The consensus binding sites of TFAP2 family members are overrepresented in lipid droplet genes. **A.** Enrichment of known transcription factor consensus sequences in lipid droplet genes. **B.** TFAP2A and TFAP2C consensus binding site motifs. **C.** Distribution of TFAP2 consensus sites in the promotor region of genes annotated to be lipid droplet related.

**Figure 2 – Figure Supplement 3.**
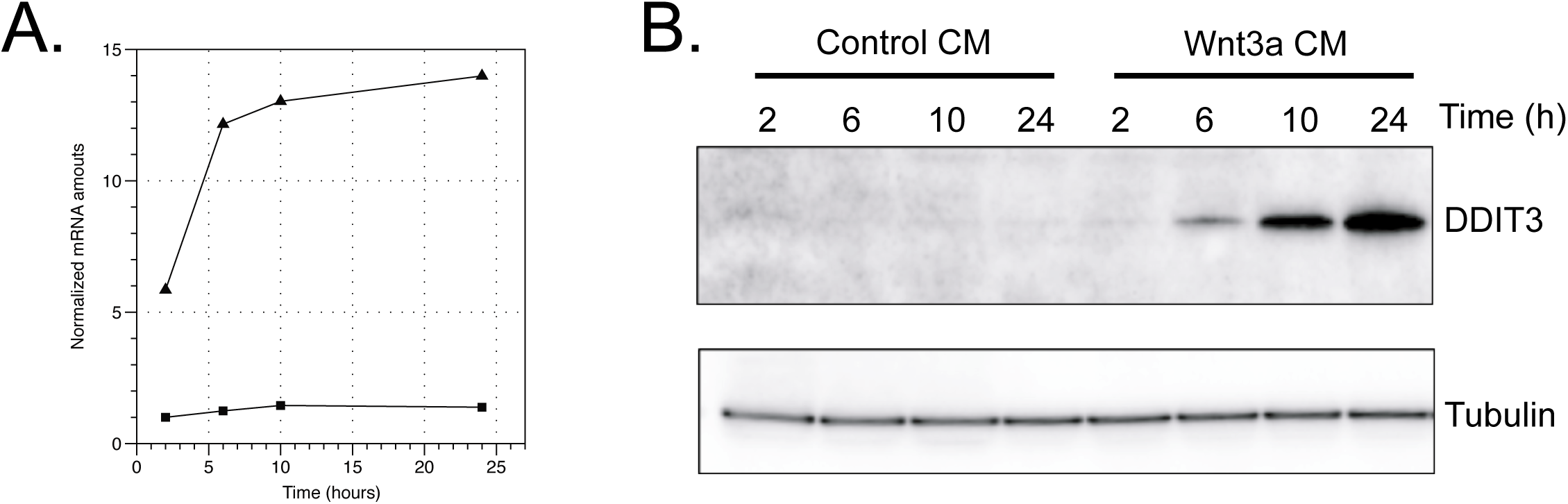
Effect of Wnt3a on DDIT3 protein and mRNA amounts in L Cells. Cells were treated with control, or Wnt3a conditioned media for the indicated times before lysis. A. mRNA was extracted from L Cells cells and amounts were determined by qPCR. B. Cell extracts were analyzed by SDS-PAGE and Western blot using antibodies against the indicated proteins.

**Figure 2 – Figure Supplement 4.**
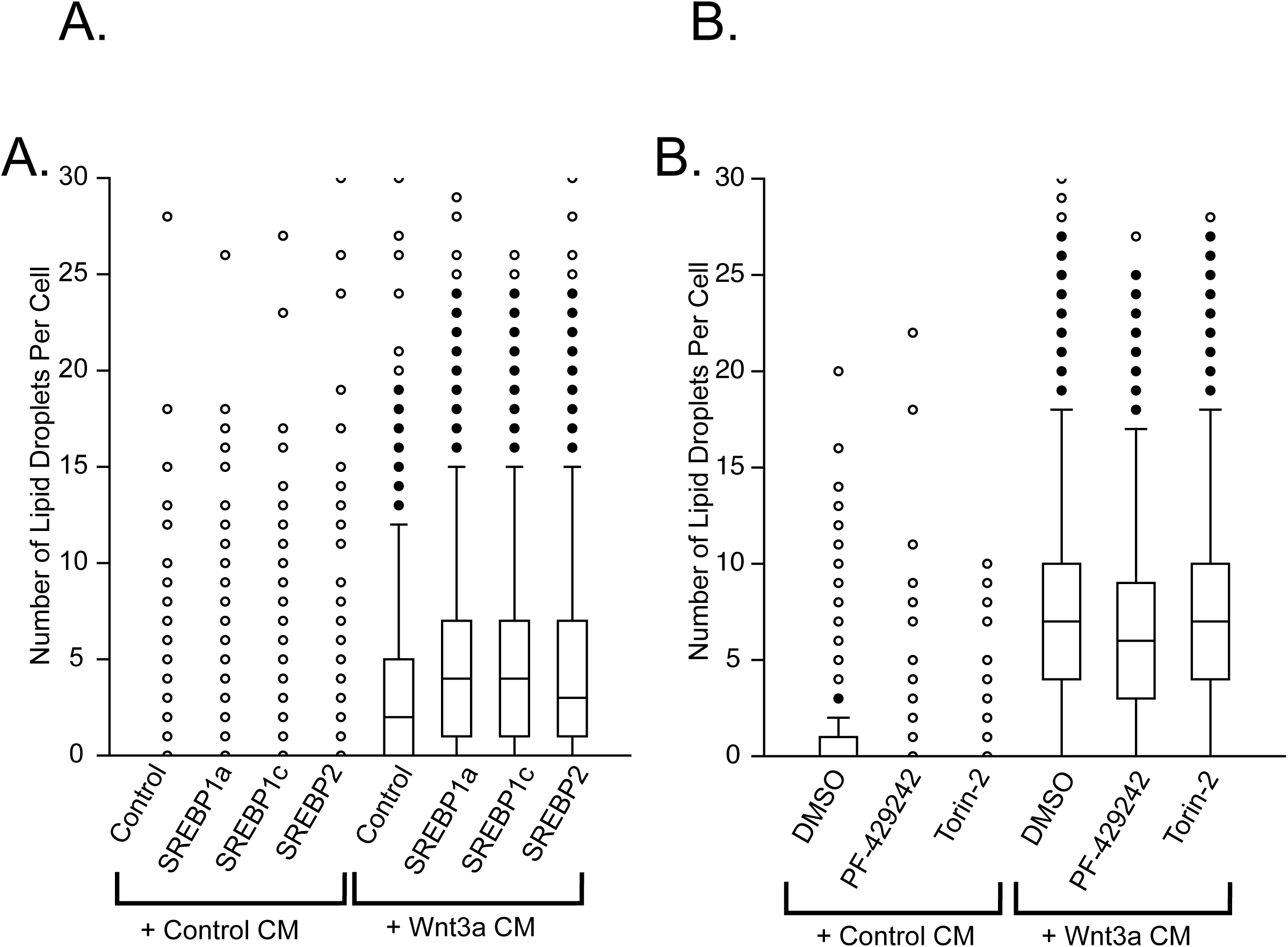
SREBF activity does not significantly influence lipid droplet number. **A.** L Cells were transfected or not with constitutively-active truncation mutants of SREBF1a, SREBF1c, and SREBF2 for 24h before addition of control or Wnt3a conditioned media for an additional 24h. Cells were fixed, labeled and imaged by automated microscopy as in Fig 1A. Data are box-and-whisker plots of a representative experiment. **B.** L Cells were treated with the indicated chemical inhibitors simultaneously with the conditioned media, and processed as (A). PF-429242; inhibitor of the SREBF site 1 protease (S1P). Torin-2; mTOR inhibitor.

**Figure 4 – Figure Supplement 1.**
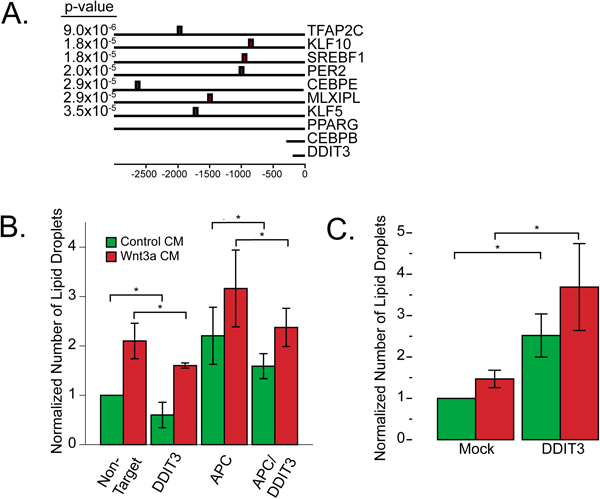
DDIT3 and lipid homeostasis. **A.** Location of DDIT3:CEBPA consensus sites in the promotor regions of the transcription factors Fig. 2B reported to be involved in lipid homeostasis. Indicated is the pvalue of the match of the consensus motif. **B.** HeLa-MZ cells were treated with siRNAs to the indicated target proteins for 48h, and then further incubated for 24h in the presence of control, or Wnt3a, conditioned media. Cells were then fixed, stained and the number of lipid droplets was quantified as in Fig 1A. Data are presented as the normalized mean number of lipid droplets per cell of 3 independent experiments ± SEM. **C.** DDIT3-mCherry was overexpressed in HeLa-MZ cells for 24h and then further incubated for 24h in the presence of control, or Wnt3a, conditioned media. Cells were then fixed and labeled, and the number of lipid droplets per cell expressing DDIT3-mCherry was quantified as in Fig 4F. Data are presented as the normalized mean number of lipid droplets per cell of 5 independent experiments ± SEM.

